# *In silico* identification of conserved *cis*-acting RNA elements in the SARS-CoV-2 genome

**DOI:** 10.1101/2020.06.23.167916

**Authors:** Bader Y. Alhatlani

## Abstract

**Aim:** The aim of this study was to computationally predict conserved RNA sequences and structures known as *cis*-acting RNA elements (CREs) located within the SARS-CoV-2 genome.

**Materials & methods:** Bioinformatics tools were used to analyse and predict *cis*-acting regulatory elements by obtaining viral sequences from available databases.

**Results:** Computational analysis prediction revealed the presence of RNA stem-loop structures within the 3’ end of the ORF1ab region that are analogous to the previously identified SARS-CoV genomic packaging signals. Alignment-based RNA secondary structures prediction of the 5’ end of the SARS-CoV-2 genome identified also conserved CREs.

**Conclusion:** These CREs could be used as potential targets for a vaccine and/or antiviral therapeutics developments; however, further studies would be required to confirm their roles in the SARS-CoV-2 life cycle.

## Introduction

In December 2019, a novel coronavirus initially named as 2019-nCoV was found to be responsible for an outbreak of pneumonia in patients in a seafood market in Wuhan, China (1). The virus was subsequently renamed as Severe Acute Respiratory Syndrome 2 (SARS-CoV-2) and identified to be from the *Betacoronavirus* genus (2). The SARS-CoV-2 is known to cause the ongoing global pandemic of coronavirus disease 2019 (COVID-19), which as of May 22, 2020 has caused more than 5 million confirmed cases with more than 332,000 deaths worldwide according to the World Health Organization (WHO). Although great efforts have been made for the development of an effective vaccine or specific antiviral treatment, there is still non available yet. The *Coronaviridae* family contains a variety of viruses that cause a wide range of diseases including respiratory, enteric, hepatic, and neurological diseases in human and animals (3). Most of the human coronaviruses (HuCoVs) usually cause mild symptoms; however, two HuCoVs which are known as Severe Acute Respiratory Syndrome (SARS-CoV) and Middle East Respiratory Syndrome (MERS-CoV), were identified to be highly pathogenic in humans (4,5). Coronaviruses (CoVs) are generally grouped into four genera: *Alphacoronavirus, Betacoronavirus, Gammacoronavirus*, and *Deltacoronavirus* (6,7).

CoVs have the largest known genomes among all RNA viruses, which range from about 26 to 32 kilobases in length, and contain enveloped positive-sense, single-stranded RNA molecule (+ve ssRNA) that is capped at the 5’ end and polyadenylated at the 3’ end (8). SARS-CoV2 shares genomic features with other SARS-like CoVs with complete genomic similarities of about 88% to bat SARS-like CoVs and 79% to the SARS-CoV (9). The SARS-CoV-2 genome is organized into about 13 open reading frames (ORFs), and two-thirds of the genome occupied by the 5’-terminal overlapping ORF1a and ORF1b which are translated from the genomic RNA to encode the replicase polyproteins pp1a and pp1b (9,10).

Like most RNA viruses, the genome of CoVs contains *cis*-acting regulatory elements (CREs) of RNA sequences and stem-loop structures that interact with RNA and viral or host proteins to form RNA-RNA or RNA-protein interactions and facilitate the viral replication, translation, and genome packaging (11,12). While often these CREs are located at the 5’ and 3’ untranslated regions (UTRs), they can also be found within the coding regions of the CoVs genome (13–15). These important regions of the viral genome maybe used as targets for the development of antiviral therapeutics for SARS-CoV-2.

## Materials and methods

### GenBank accession numbers of viral sequences

Viral genomic sequences were retrieved from the GenBank in the National Center for Biotechnology Information (NCBI). Virus strains and accession ID used in this study are as follow: (NC_045512.2) for SARS-CoV-2, (NC_004718.3) for SARS-CoV, and (MG772933) for Bat SARS-like CoV.

### Bioinformatics analysis

The RNA secondary structures of the viral genomic sequences were predicted using the online Mfold Web server at http://unafold.rna.albany.edu/?q=mfold/, and RNAfold Web server at http://rna.tbi.univie.ac.at/cgi-bin/RNAWebSuite/RNAfold.cgi (16,17). In addition, LocARNA Web server at http://rna.informatik.uni-freiburg.de/LocARNA/Input.jsp was used for the alignment-based prediction of consensus RNA secondary structures at the 5’-terminal region of SARS-CoV-2, bat SARS-like CoV, and SARS-CoV genomes (18). The VARNA Web applet at http://varna.lri.fr/ was then used to draw the RNA secondary structures (19).

## Results

Previous studies on some betacoronaviruses such as MHV, MERS-CoV, and SARS-CoV revealed that the ORF1b region may contain CREs that are suggested to function as packaging signals (PSs) (20–23). These CREs are functionally and structurally conserved within the same lineages of betacoronavirus (14). A previous bioinformatics study on the SARS-CoV genome predicted a stable stem-loop RNA structure at the 3’ end of ORF1b, encompassing nucleotides (ntds) 19888 to 19950, to be the putative core PS (PS_core_) of SARS-CoV (22). A further functional analysis identified the functional PS of the SARS-CoV as a 580 ntds encompassing the viral genomic RNA from ntds 19712 to 20294, which fold into a RNA secondary structure including the PS_core_ and binds to the neucleocapsid (N) protein (23). SARS-CoV-2 shares 87.6% and 79% complete genome similarities with bat SARS-like CoV and SARS-CoV, respectively (9,24).

Hence, in this study bioinformatics analysis was used to identify if the ORF1b region of SARS-CoV-2 pocesses such *cis*-acting RNA elements similar to that observed in SARS-CoV and other CoVs. To test this hypothesis, genome sequences of the SARS-CoV-2 ORF1b region were analyzed for RNA secondary structures and compared with the closely related SARS-CoV and bat SARS-like CoV sequences. RNA sequences of SARS-CoV-2 spanning ntds 15,000 to 21,541 were first analyzed using RNAfold Web server to predict the minimum free energy (MFE) (17). Results predicted the lowest free energy located between ntds 19,000 to 20,300 of the SARS-CoV-2 ORF1b region (Figure 1). Because this position overlaps with the genomic PS of the SARS-CoV, which was identified at position ntds 19712 to 20294, the Mfold Web server was then used to predict the secondary structures of the RNA sequences spanning ntds 19,712 to 20,294 of the SARS-CoV-2 ORF1b region and were compared with those of SARS-CoV and bat SARS-like CoV (16). The Mfold analysis resulted in differences in the predicted RNA secondary structures between the three sequences; however, two stable stem-loops were identified to be similar and observed in all of the three sequences of the viruses (Figure 2A-C). The predicted two stem-loops (named as SL1 and SL2) located at position 19,900 – 20,000, 19,839 – 19,943, and 19,903 – 20,000 of the SARS-CoV-2, SARS-CoV, and bat SARS-like CoV genomes, respectively (Figure 3A-C). The upper part of SL1 which contains 38 ntds is structurally conserved between the three viruses with covariation in the sequences. In the case of SARS-CoV-2, the SL1 is longer than those predicted in SARS-CoV and bat SARS-like CoV with 70 ntds in length, whereas its length in the SARS-CoV is 51 ntds, and only the upper part of SL1 is formed in bat SARS-like CoV (Figure 3A-C). The SL2 is much shorter in SARS-CoV-2 with only 26 ntds in length compared to that predicted in SARS-CoV and bat SARS-like CoV which contains 51 ntds in both viruses. However, the genetic sequences of the upper part of SL2 is more conserved among the three viruses than SL1, with only two nucleotides differences between SARS-CoV-2, SARS-CoV and bat SARS-like CoV. The first covariant is located at the stem and includes a C-G base pair in the SARS-CoV-2 sequence in place of a U-G base pair found in SARS-CoV and bat SARS-like CoV (Figure 3A-C). This single nucleotide difference from a C to U maintains the base pairing of the stem and does not change the amino acid (Leu) of ORF1b (Figure 3A-C). The second covariant is located at the U-rich loop, and it is the third base in the loop in which this nucleotide is a U in the SL2 of SARS-CoV-2, whereas it is a G and an A in the SARS-CoV and bat SARS-like CoV sequences, respectively. However, this single nucleotide difference changed the amino acid from (Leu) in the case of SARS-CoV and bat SARS-like CoV to (Phe) in the SARS-CoV-2 genome (Figure 3A-C). It should be noted that the SL2 that was the previously predicted to function as the putative core PS (PS_core_) of SARS-CoV genome (22,23). Therefore, it is reasonable to assume that the predicted RNA stem-loop structures of SARS-CoV-2 may also have the same role as a putative genomic packaging signal, given the conservations of the RNA structures and sequences of the SL1 and SL2 of the viruses as well as the similarities of the genomic location of these predicted RNA structures within the ORF1b.

**Figure 1.**
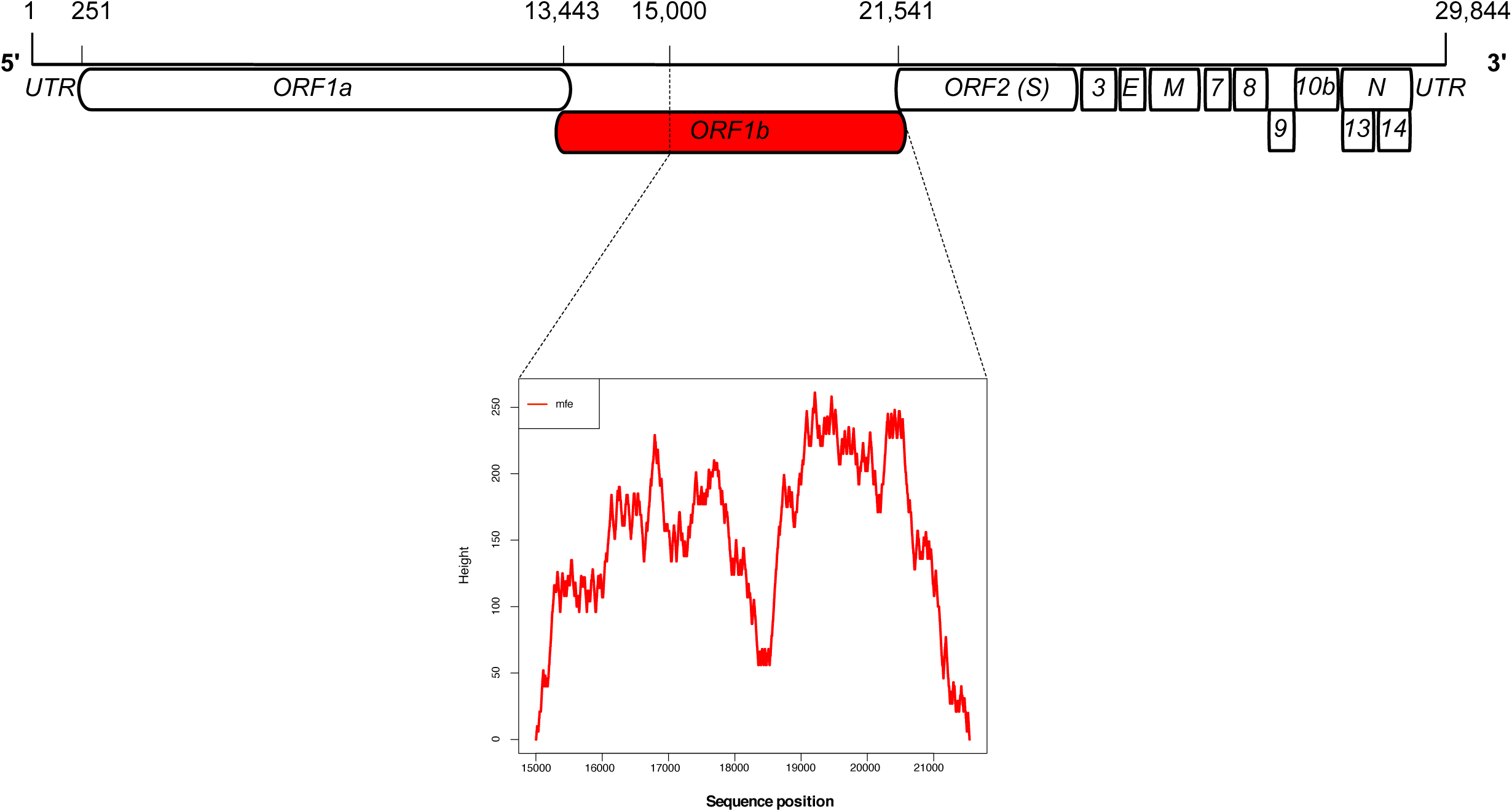
Prediction of minimum free energy (MFE) structures within the SARS-CoV-2 ORF1b region. The SARS-CoV-2 genome is about 30kb in length that is organized in thirteen open reading frames (ORFs). The viral genome is flanked by the 5’ and 3’ untranslated regions (UTRs). The ORF1b region is highlighted in red (upper panel), and a mounting plot represents prediction of the MFE structures spanning nt 15,000 to 21,541 of the SARS-CoV-2 genome sequences, (NC_045512.2) is shown with the highest peak of MFE prediction located between nt 19,000 to about 20,300 of the SARS-CoV-2 ORF1b region (lower panel). Note that the mountain plot indicates the secondary structure in a plot of height versus sequence position, where the peaks represent the probabilities of RNA base pairing.

**Figure 2.**
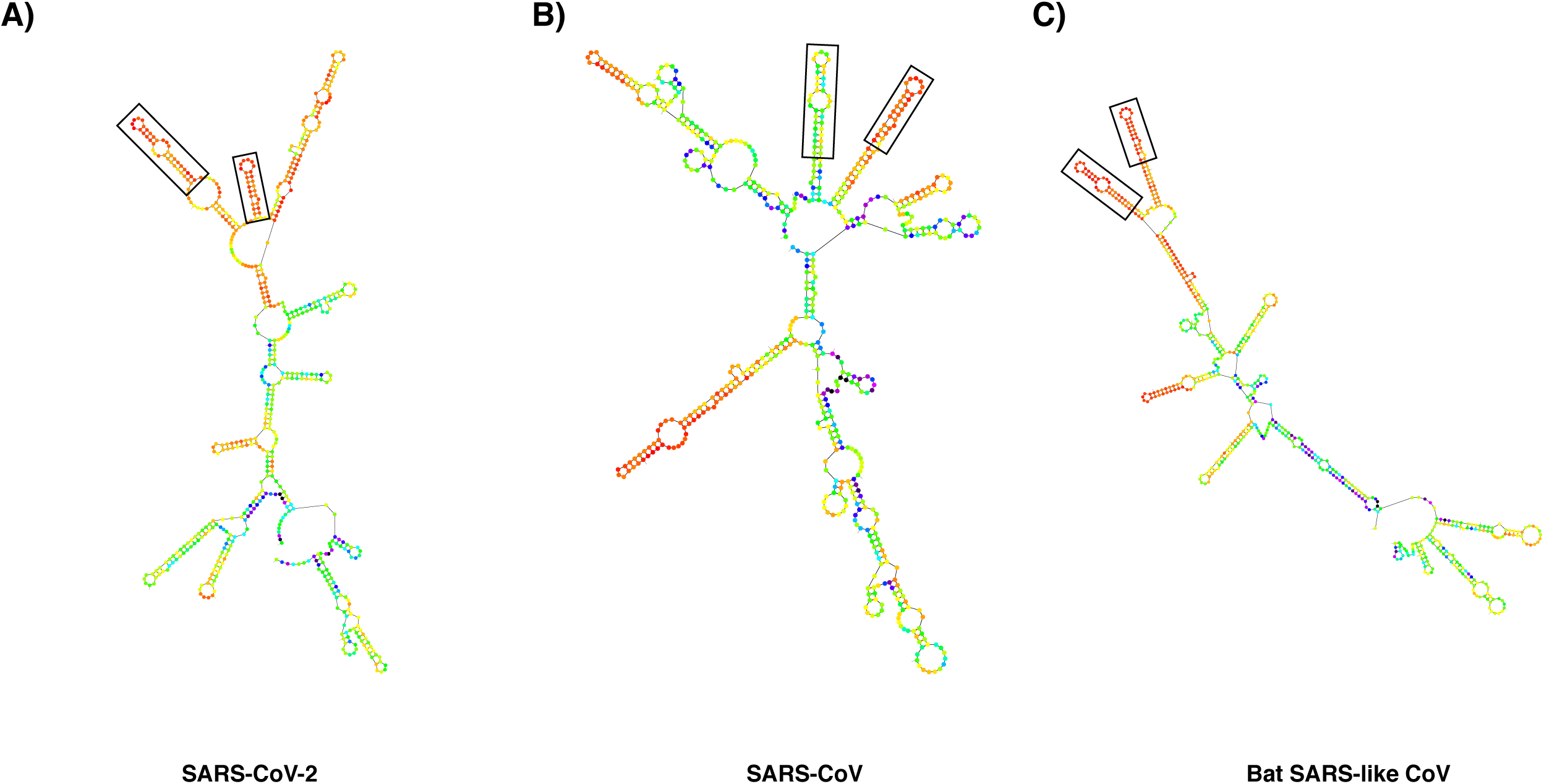
Prediction of the ORF1b RNA secondary structures. RNA secondary structures located at the 3’ end of the ORF1b region of SARS-CoV-2 (nt 19,712 to 20,294) was predicted using the Mfold Web server (A) and compared with RNA secondary structures at the same position of SARS-CoV (NC_004718.3) (B) and bat SARS-like CoV (MG772933) (C). The two identical RNA stem-loops (SL1 and SL2) are indicated in rectangle in the three viruses.

**Figure 3.**
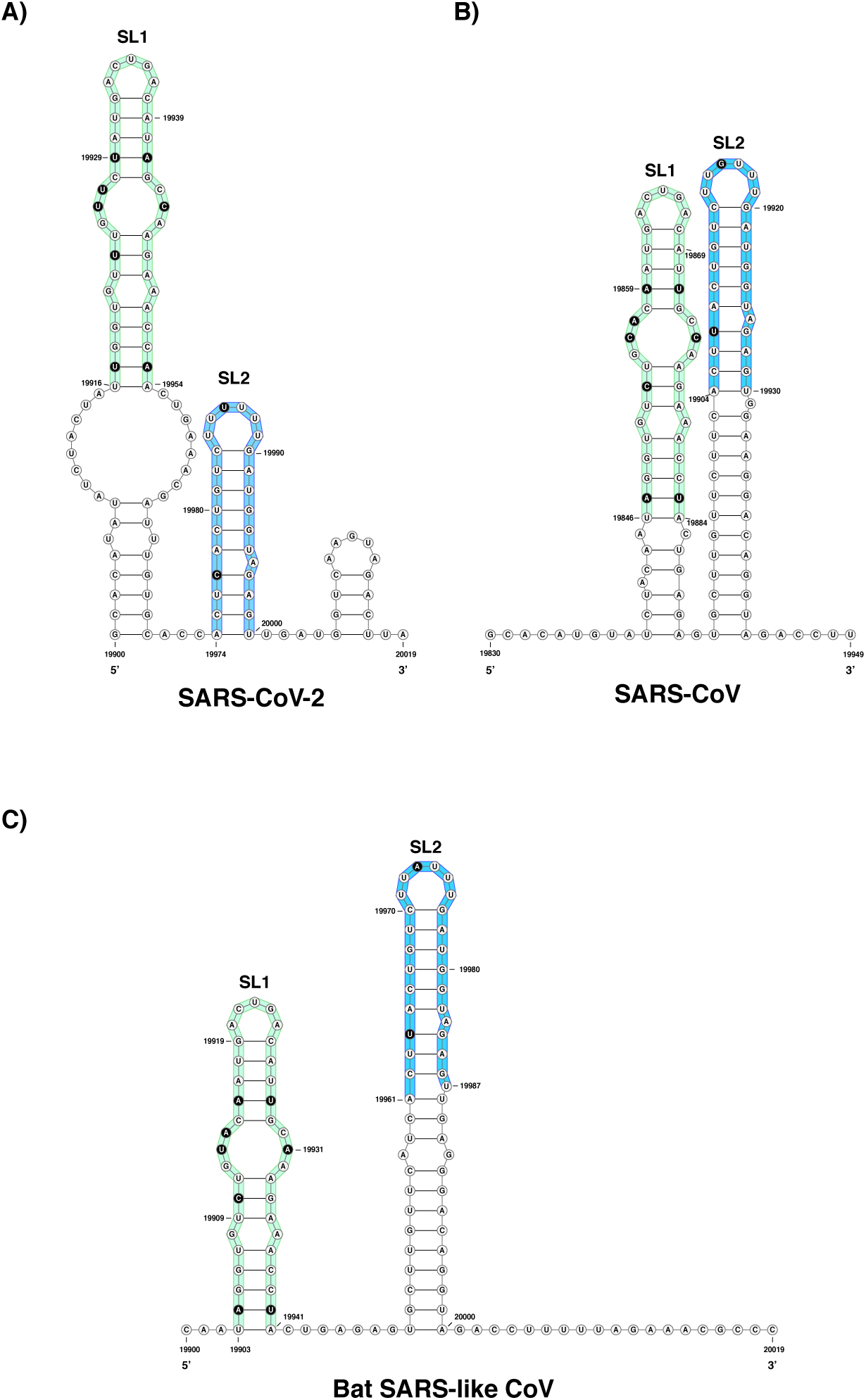
Comparison of the predicted RNA secondary structures among SARS-CoVs. The viral RNA sequences in the 3’ end of ORF1b region from 19,900 to 20,019 for both SARS-CoV-2 and bat SARS-like CoV, and in the region of 19,830 to 19,949 for SARS-CoV, were folded using the Mfold Web server and drawn using the VARNA Web applet to compare between the predicted RNA stem-loops. The similarities of the RNA structures of the predicted SL1 and SL2 between the three viruses are highlighted in green and blue, respectively. In addition, the sequence variation in these two stem-loops are indicated in black. Note that these two RNA stem-loops are part of the previously identified SARS-CoV genomic PS, with SL2 being identified as the PS_core_ (22,23).

*Cis*-acting regulatory elements of RNA secondary structures and sequences have also been previously described at the 5’ end of a number of CoVs including members of the *Betacoronavirus* genus such as mouse hepatitis coronavirus (MHV), bovine coronavirus (BCoV), MERS-CoV, and SARS-CoV (15). The model of the RNA secondary structures at the 5’ region of the SARS-CoV genome was previously predicted to fold into eight stem-loops (SL1 to SL8) (15). To predict the RNA secondary structures at the 5’-proximal sequence of the SARS-CoV-2 genome and compare it to that previously identified in the 5’ end of the SARS-CoV genome and also to the closely related bat SARS-like CoV, the first 474 ntds of the SARS-CoV-2 genomic sequence was analysed using the Mfold Web server. It should be noted that during the preparation of this manuscript, a recent study has predicted some conserved RNA structures within the SARS-CoV-2 genome including those RNA elements at the 5’ and 3’ ends of the viral genome (25). Results of the Mfold analysis predicted an identical 5’-terminal RNA secondary structures model of the SARS-CoV-2 to the SARS-CoV model which was previously proposed by Yang et al., with eight RNA stem-loops (SL1 to SL8) (Figure 4). In addition, the predicted model in this study is similar to that recently described in the bioinformatics study by Rangan et al., except that they have predicted seven stem-loops (SL1 to SL7), while in this study an additional stem-loop (SL8) was identified which is consistent with the model for SARS-CoV 5’ RNA structure (15,25). The SL1, SL2, and SL4 are located within the 5’ UTRs of coronavirus genomes and they are structurally conserved among at least three genera of coronaviruses, with the SL2 being the most conserved RNA secondary structure at the 5’ UTR of coronavirus genomes (26). In addition, results also demonstrated that the conserved core leader of the transcriptional regulatory sequence (TRS-L) region required for subgenomic RNA (sgRNA) synthesis is located within the SL3 which is consistent with previous studies of SARS-CoV and BCoV (Figure 4) (26). The predicted SL4, which has been shown to be conserved in all CoVs, is longer than the three preceding stem-loops and contains a short upstream ORF (uORF) that is found in most CoVs (13,26,27). Moreover, a long stem-loop RNA structure that contains three hairpin substructures (termed as SL5A, 5B, and 5C) was also identified (Figure 4) (14). Part of the SL5 is located in the 5’ UTR; however, the AUG initiation codon of the nonstructural protein 1 (nsp1) is located downstream of SL5C at a position similar with previous studies on SARS-CoV (Figure 4). It should be noted that the SL5ABC is conserved among betacoronaviruses, suggesting the essential roles of these stem-loops structures as *cis*-acting regulatory elements in the life cycle of CoVs such as viral replication. In agreement with a previous study on SARS-CoV, the loops of the SL5A and SL5B contain the conserved 5’-UUUCGU-3’ motifs, and this is equivalent to the conserved 5’-UUYCGU-3’ loop sequences that was found in the SL5ABC of alphacoronaviruses (13,26). Bioinformatics analysis predicted also three stem-loops (SL6, SL7, and SL8) in the nsp1 coding sequence similar to that found in the 5’-terminal of the SARS-CoV (Figure 4). However, these RNA structures are known to be less conserved between CoVs lineages than the stem-loops found within the 5’ UTR of CoVs genomes.

**Figure 4.**
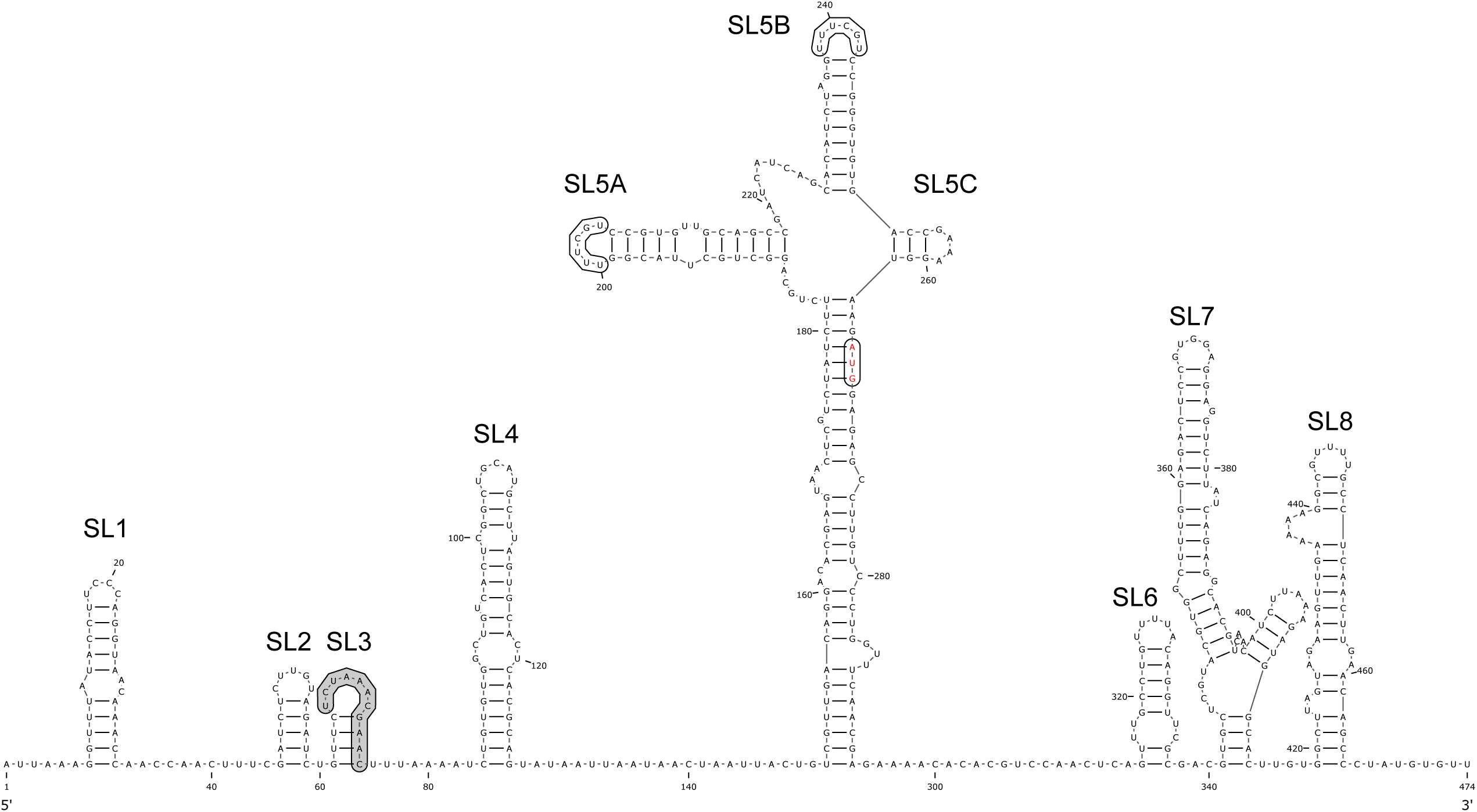
Computational prediction of RNA secondary structures model at the 5’-terminal of the SARS-CoV-2 genome. Schematic representation of the RNA secondary structures at the 5’ region of the SARS-CoV-2 genome (nt 1 to 474) was generated using the Mfold Web server and drawn using the VARNA Web applet (ΔG = −155.50 kcal/mol). The eight RNA stem-loops (SL1 to SL8) are indicated, and the conserved core leader TRS region located within the SL3 is highlighted in grey. In addition, the conserved 5’-UUUCGU-3’ motifs found within the loop of SL5A and SL5B is indicated in boxes, and the AUG that represents the start codon of the ORF1ab is indicated in red. The SARS-CoV-2 reference genome sequence obtained from GenBank (NC_045512.2) was used.

To further confirm the conservation of the RNA secondary structures located at the 5’ region of the viral genome among SARS-related viruses, the first 474 ntds of SARS-CoV-2, SARS-CoV, and bat SARS-like CoV genomes were aligned and folded using LocARNA Web server (18). Results from sequences alignments indicated that all the RNA stem-loops are highly conserved among the three viruses, with SL2 and SL3 being the most conserved among the other stem-loops which have sequences covariation (Figure 5). This high degree of conservation is expected for SL2 and SL3 because as mentioned above that SL2 is suggested to be the most conserved RNA elements in the CoVs 5’ UTR region, and SL3, which is found in SARS-like CoVs and BCoVs, contains the TRS-L sequences that have an essential role in the synthesis of sgRNA (13,28). These results indicated the high conservation of the RNA elements at the 5’ end of the SARS-like CoVs genomes, and hence suggesting that these RNA secondary structures also function as *cis*-acting regulatory elements in the life cycle of SARS-CoV-2.

**Figure 5.**
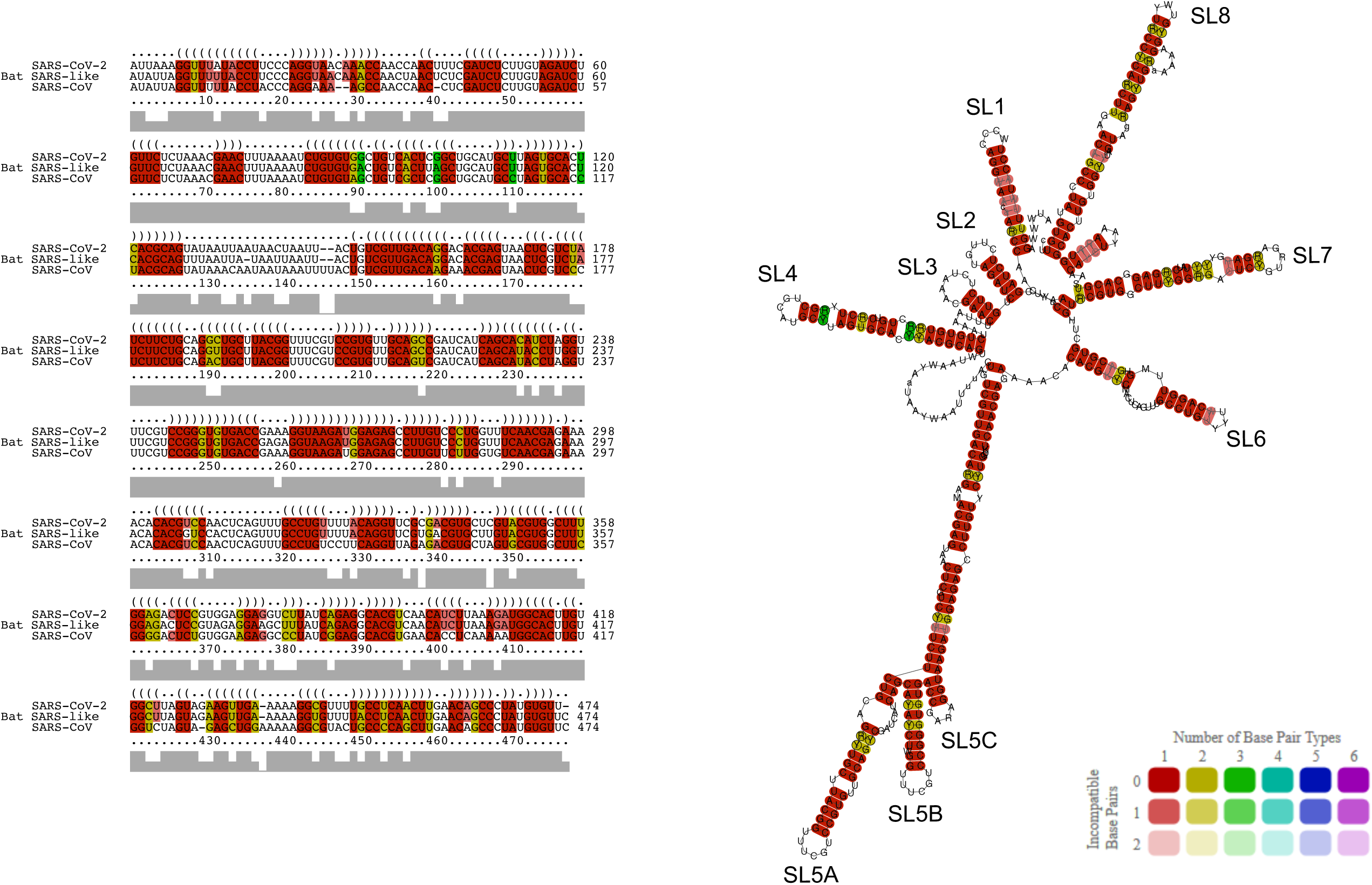
Alignment-based prediction of the RNA structures at the 5’-terminal of SARS-CoVs genomes. The first 474 ntds of the SARS-CoV-2, bat SARS-like CoV, and SARS-CoV viral genomes were aligned using LocARNA to predict conserved RNA stem-loops at the 5’ end of the viral genomes. Colours indicate the conservation of base pairs types where red represents conservation of one base pairs and yellow and green indicate two and three base pairs, respectively. Dark colour represents that all the three virus contain this bas-pair, whereas light colour shows that one or two of the sequences do not contain the base pairs.

## Discussion

*Cis*-acting regulatory elements have been described and characterized in the genomes of several RNA viruses, including picornaviruses, coronaviruses, and noroviruses (11,29,30). These CREs play important functional roles in the viral life cycle and usually contribute to the synthesis of viral RNA replication, translation, and genome packaging (11). Previous studies have identified stem-loop RNA structures, located at the 3’ end of the SARS-CoV ORF1b, which are recognized by the SARS-CoV structural N protein and suggested to act as the genomic PS (22,23). Therefore, this work aimed to use bioinformatics tools in order to identify if the genome of the newly emerged SARS-CoV-2 shares conserved CREs that are present within the 5’ end and the ORF1ab regions of the viral genome.

Computational analysis predicted two conserved stem-loops within the 3’ end of the ORF1b region (named as SL1 and SL2), the latter of which have been previously described to be part of the SARS-CoV PS and was named as (PS_core_) (22). The SL2 is more conserved between the three viruses than SL1, and the top of SL2 is featured with a hexaloop that contains a U-rich motif. It should be noted that genomic PSs for different CoVs consist of RNA structure elements that vary in length and location in the genome of viruses within the same lineages (14,31). For example, the genomic PS of MERS-CoV which is a lineage C betacoronavirus was identified in a similar position as the SARS-CoV at the 3’ end of the ORF1ab region (21). Moreover, the genomic PS is functionally and structurally conserved in lineage A betacoronaviruses which contains a 95 ntds stem-loop RNA structure and located within the 3’ region of ORF1b (32). However, the genomic PS of transmissible gastroenteritis virus (TGEV) is located at the 5’-terminal of the viral genome (33). Although this study cannot conclude the functionality of the predicted RNA stem-loop structures located at the SARS-CoV-2 ORF1b region and their roles in the SARS-CoV-2 life cycle, it is postulated that they could be functioning similarly as *cis*-acting elements and hence they could be a putative genomic PS for the SARS-CoV-2. This is because of (i) the similar position of the viral genome where these RNA elements locate at the 3’ end of ORF1b, (ii) and also because of the conserved RNA secondary structures and sequences of these predicted stem-loops when compared to those found in the closely related SARS-CoV and bat SARS-like CoV.

This study has also used genome sequences at the 5’-proximal region of the SARS-CoV-2 genome to predict consensus conserved RNA secondary structures by comparing with SARS-related CoVs, such as SARS-CoV and bat SARS-like CoV. The predicted model in this study was similar to a previous model that was described for SARS-CoV, with eight RNA stem-loops (SL1-SL8) were identified (15). In addition, this model of RNA structures at the 5’ region of SARS-CoV-2 is similar to a model recently described in a bioinformatics study with only difference in the SL8 which was not included in that study (25). It has been suggested that the RNA secondary structures of SL1, SL2, and SL4 are conserved in three genera of CoVs, whereas SL1, SL2, and SL4 and SL5ABC are conserved among betacoronaviruses only (15).

## Conclusion

In summary, this study used computational tools to predict *cis*-acting RNA motifs that present at the SARS-CoV-2 RNA genome. Bioinformatics analysis suggested that the 3’ end of the SARS-CoV-2 ORF1b region may contain RNA structure elements that are structurally conserved in other SARS-CoVs, and analogous to the SARS-CoV genomic packaging signal. In addition, this study also demonstrated and confirmed the predicted RNA secondary structures at the 5’-proximal region of the SARS-CoV-2 genome. However, further studies would be required to (i) biochemically confirm these *cis*-acting RNA elements, (ii) and to further investigate their roles in the viral life cycles. These important regions within the viral genome could be then targeted for the development of a vaccine and/or antiviral therapeutics against this highly important pathogen.

